# Glycolytic oscillations under periodic drivings

**DOI:** 10.1101/2023.10.08.561464

**Authors:** Pureun Kim, Changbong Hyeon

**Affiliations:** Korea Institute for Advanced Study, Seoul 02455, Korea

## Abstract

In many living organisms displaying circadian rhythms, the intake of energy often occurs in a periodic manner. Glycolysis is a prototypical biochemical reaction that exhibits a self-sustained oscillation under continuous injection of glucose. Here we study the effect of periodic injection of glucose on the glycolytic oscillation from a dynamical systems perspective. In particular, we employ the Goldbeter’s allosteric model of phosphofructokinase (PFK) as a model system for glycolytic oscillations, and explore the effect of periodic substrate influx of varying frequencies and amplitudes by building the phase diagrams of Lyapunov exponents and oscillatory periods. When the frequency of driving is tuned around the harmonic and sub/super-harmonic conditions of the natural frequency, the system is entrained to a frequency-locked state, forming an entrainment band that broadens with an increasing amplitude of driving. On the other hand, if the amplitude is substantial, the system may transition, albeit infrequent, to a chaotic state which defies prediction of dynamical behavior. Our study offers in-depth understandings into the controllability of glycolytic oscillation as well as explains physical underpinnings that enable the synchronous oscillations among a dense population of cells.

---

Glycolysis is one of the metabolic processes essential for all living organisms. It is an anaerobic process that catalyzes the transformation of one glucose molecule into two pyruvates, each resulting in a net production of 2 ATPs and a reduction of NAD^+^ to NADH, which are later used in the processes of Kreb cycle and electron transfer cycle to generate more ATPs under aerobic condition [1]. Notably, in yeast and heart muscle cell extracts [2–4] and in pancreatic *β*-cells [5–7], the products of glycolysis exhibit sustained oscillations even when the influx of glucose to the system is constantly maintained.

Among various models and mechanisms developed for describing glycolytic oscillations [8–12], the positively regulated autocatalytic-allosteric action of phosphofructokinase (PFK) has been suggested as a core mechanism for the generation of the sustained oscillation [4, 8, 13, 14]. Specifically, PFK is a phosphotransferase that catalyzes ATP hydrolysis, so as to convert fructose 6-phosphate (F6P) into fructose 1,6-biphosphate (FBP). Binding of one of the reaction products (ADP) to the regulatory sites of PFK positively regulates the allosteric transition of PFK into a catalytically more competent state, which generates oscillations in substrate and product concentrations. In experiments using yeast cell extracts, sustained oscillations of NADH concentration were observed as long as the influx of glucose and the degradation of product were maintained in a proper range [4, 15].

From the perspective of dynamical systems of continuous variables in two dimensions, the sustained oscillation of substrate and product concentrations ([*S*] and [*P* ]) can be understood as a stable limit cycle formed around an unstable fixed point, which is ensured by the Poincaré-Bendixon theorem [16]. An introduction of third variable that changes over time, such as a non-autonomous time-varying influx of glucose to the system, however, substantially increases the complexity of the resulting dynamics. In particular, the external periodic driving effectively puts dynamical trajectories on the phase space surface of a torus [17, 18]. Boiteux *et al*. [4] carried out an experimental study to explore the effect of stochastic and *periodic* injection of glucose on the glycolytic oscillation, and observed subharmonic entrainments of the system for the case of periodic injection (see Appendix for the effect of stochastic forcing). That is, when the period of external driving (*T*_ext_) relative to the autonomous period of the system (*T*_*o*_) was over a certain range, the system was entrained towards a new period of *T*_obs_ = *nT*_ext_ (*n* = 1, 2, 3, …). Meanwhile, if the system was not entrained, the external driving gave rise to an irregular trajectory.

Our study on the PFK model under periodic influx of substrate can be considered as a theoretical extension of the Boiteux *et al*.’s experimental work [4]. We find that the perturbation leads to extremely rich dynamical behaviors. By varying the frequency (*ω*_ext_ = 2*π/T*_ext_) and amplitudes (*ε*) of the external driving, we calculate 2D phase diagrams of (i) the Lyapunov exponent (*λ*) and (ii) the resulting oscillation period (*T*_obs_), which not only clarify the condition that leads to entrainment, but also help differentiate between chaotic and quasi-periodicity orbits. Our study offers in-depth understandings into the controllability of glycolytic oscillation by illuminating its dynamical responses and modulations under periodic drivings.

## PFK MODEL OF GLYCOLYTIC OSCILLATION

The PFK is an oligomeric enzyme that retains a catalytic and a regulatory site in each of *n* subunits, where the value of *n*(= 2, 4, 8) varies with the species. At low substrate concentration, the PFK is predominantly in the tense (*T* ) state, but high concentration of substrate shifts the pre-equilibrated population of enzyme in *T* state to the *catalytically competent* relaxed (*R*) state. The PFK in the *R* state is capable of converting F6P into FBP via phosphorylation while catalyzing the ATP hydrolysis. A reaction product, ADP, cooperatively binds to the *n* (= 8 for yeast) distinct regulatory sites available in the *R* state and positively regulates the *T → R* transition of PFK. Thus, with an increasing substrate concentration, PFK has more chances to exhibit its catalytic power, which accelerates with the amount of ADP until the *n* regulatory sites are half filled by ADP but saturates when all the sites are filled up, giving rise to an activity curve with a sigmoidal shape.

The autocatalytic-allosteric model of glycolysis processed by PFK is formulated using a system of ordinary differential equations (ODEs) [8, 19]:

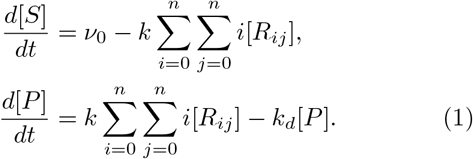

The ODEs describe the mass balance of the substrate and product concentrations. The substrate *S* supplied to the system with a rate *ν*_0_ is converted by PFK to a product *P* with a rate proportional to the occupation number of the substrate in the catalytic sites of *R* state, 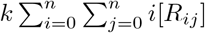 where *k* denotes the conversion rate and *R*_*ij*_ denotes a microstate of *R* state in which *i*(= 0, 1, …, *n*) catalytic sites and *j*(= 0, 1, …, *n*) regulatory sites are occupied by the substrate and product, respectively. The product *P* undergoes uni-molecular degradation with the rate *k*_*d*_ from the system. In writing the ODEs in Eq. 1, it is assumed that the conformational changes of PFK enzyme into its subpopulations of *T* or *R* states occur in much faster time scales than the time scale associated with the variations in substrate and product concentration. As a consequence, the corresponding concentrations of the subpopulations [*T*_*i*_] and [*R*_*ij*_] are pre-equilibrated and can be expressed as a function of the substrate and product concentration ([*S*] and [*P* ]) (see Eqs. 2 and 3). The PFK model, proposed by Goldbeter [8], is built based on the Monod-Wyman-Changeux (MWC) model [20–22], which is the workhorse of protein allostery [22–25], along with a set of key parameters:

- *L* ≡ [*T*_0_]*/*[*R*_0_], where *R*_0_ ≡ *R*_00_, is the ratio of *T* and *R* states in the absence of substrate.
- 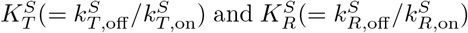 are the dissociation constants or the binding affinities of substrate (*S*) to the *T* and *R* states, respectively.
- 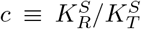, the ratio of binding affinities in *R* and *T* states, specifies the preference of substrate binding to the *R* against the *T* state.

In the PFK model, the concentration of PFK subpopulation in the *T* state bound by *i* substrates (0 ≤ *i* ≤ *n*) can be expressed as

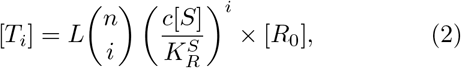

and similarly the concentration of PFK subpopulation in the *R* state bound by *i* substrates in the catalytic sites and *j* product molecules in the regulatory sites is given as

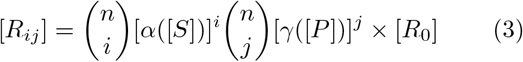

where 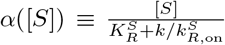 and 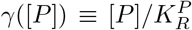 with 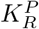 denoting the binding affinity of product (ADP) to the regulatory site of the *R* state. Using Eqs.2, 3 and the total enzyme concentration 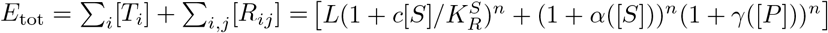 [*R*_0_], one can express 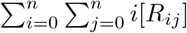 in terms of [*S*] and [*P* ] as

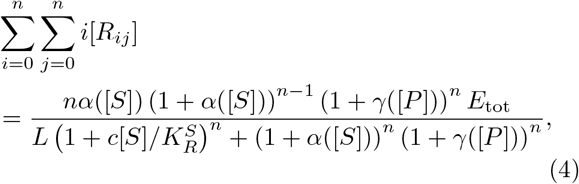

which clarifies that Eq. 1 is a set of ODEs with two variables, 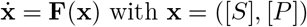.

In our earlier work [26], we used the parameters, 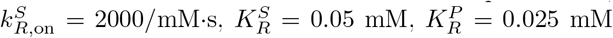, [27–30], and 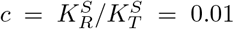 which ensures pref-erential binding of substrate to the *R* state. Since the ATP hydrolysis time due to ATPase activity is typically ≳ 𝒪 (1) msec [31], we used *k* = 500 s^−1^ for the rate of catalysis. In addition, by specifically considering the yeast PFK that adopts an octameric form (*n* = 8), we use the allosteric constant *L* = 4 × 10^9^ which is 3 orders of magnitude greater than the value known for the tetramer (*n* = 4) [29]. In Ref. [26], we studied the phase diagrams of various quantities as a function of constant influx of substrate *ν*_0_ and the degradation rate *k*_*d*_, identifying the parameter space associated with the self-sustained oscillations (Fig. 1B). For the parameter values relevant for octameric PFK of yeast (*L* = 4 × 10^9^, *k* = 500 *s*^−1^, *ν*_0_ = 0.005 mM·*s*^−1^, *k*_*d*_ = 0.05 *s*^−1^, *c* = 0.01, *E*_tot_ = 5 × 10^−4^ mM, and *n* = 8), which yields a stable oscillation with a period of *T*_*o*_ = 388.6 s, we find the system poised at a thermodynamically optimal point with minimal entropy production rate [26, 32] in the limit cycle region near the phase boundary (the point marked with the yellow star in Fig. 1B), corresponding to the state above the Hopf bifurcation point [26].

**FIG. 1.**
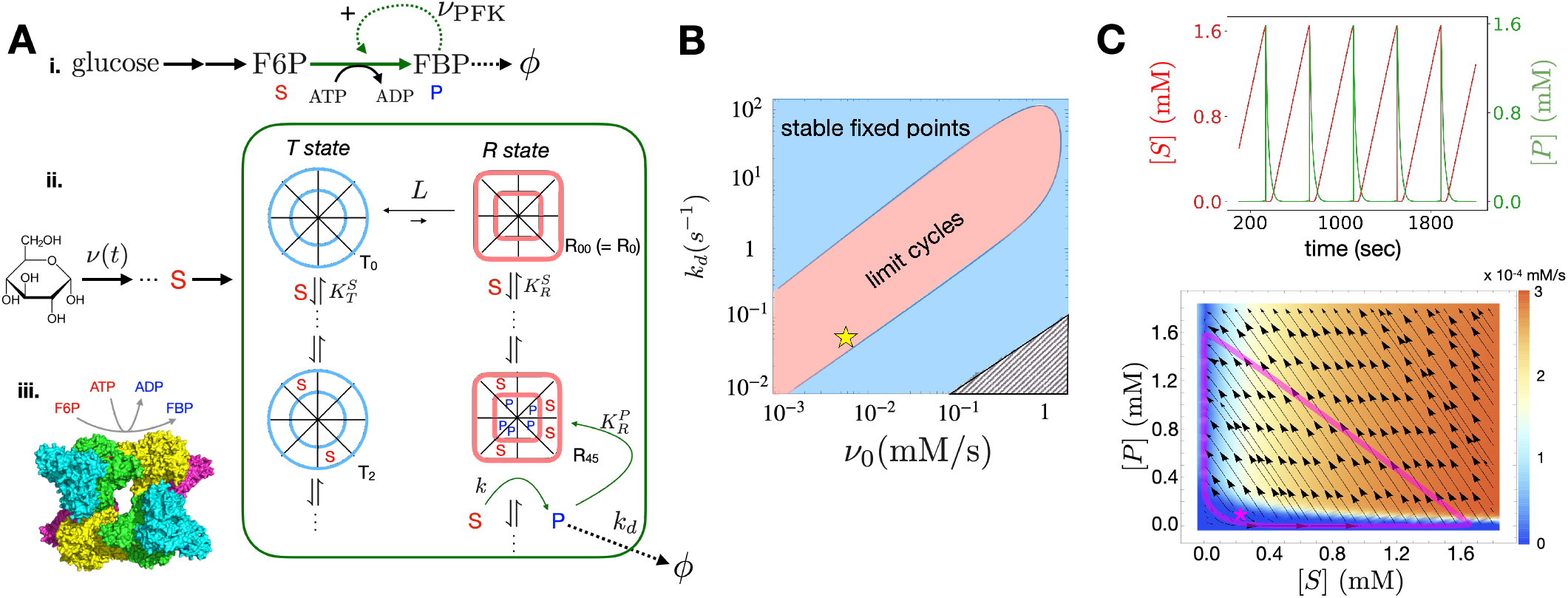
Allosteric model for phosphofructokinase. (A) (i) The schematic of glycolytic pathway, highlighting the F6P FBP conversion regulated by PFK that undergoes positive allosteric activation. (ii) The more detailed schematic highlighting the allosterism of PFK catalysis. Each subunit of PFK has catalytic and regulatory sites, where substrate (*S* = F6P) and product (*P* = ADP) bind, respectively. The oscillatory dynamics of PFK model is realized in a setting of open thermodynamic system, where glucose molecules, which leads to the production of F6P, are injected at the rate *ν* (*t*) and the product (*P* ) degrades at the rate of *k*_*d*_. At low substrate concentration 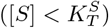, the enzyme is mainly in the *T* state. But, with increasing substrate concentration, conformational transition to the *R* state occurs, increasing the chance of enzymatic activity. (iii) The octameric structure (*n* = 8) of PFK (left bottom corner), which catalyzes the conversion of F6P to FBP via the ATP hydrolysis. (B) Phase diagram of dynamical behavior of PFK model as a function of constant injection rate *ν* (*t*) = *ν* _0_ and degradation rate *k*_*d*_. A set of parameter values relevant for octameric PFK of yeast (*L* = 4 × 10^9^, *k* = 500 *s*^−1^, *ν*_0_ = 0.005 mM · *s*^−1^, *k*_*d*_ = 0.05 *s*^−1^, *c* = 0.01, *E*_tot_ = 5 × 10^−4^ mM, and *n* = 8), which corresponds to the location marked with a yellow star on the phase diagram, yields a stable limit cycle with a period of *T*_*o*_ = 388.6 sec. (C) Anti-phasic oscillations of [*S*](*t*) and [*P* ](*t*) (top), and the phase portrait (bottom) with the velocity fields 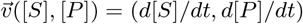 along with the steady state trajectory (thick magenta line) around the unstable fixed point ([*S*^*^], [*P* ^*^]) = (0.22 mM, 0.10 mM) marked by an asterisk. The color-code in the phase portrait depicts the size of the velocity field, 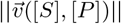.

Here, we explore how the oscillatory behavior of the PFK model is modulated under a periodic influx of substrate *ν* (*t*) satisfying

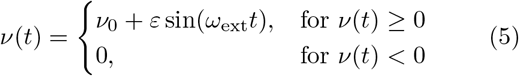

with *ω*_ext_ = 2*π/T*_ext_. For *ε > ν* _0_, *ν* (*t*) is set to 0 if *ν* (*t*) *<* 0, which renders *ν* (*t*) a strong periodic pulse to the system.

The main set of ODEs, Eq. 1 with Eq. 4, was numerically integrated to generate time trajectories of [*S*](*t*) and [*P* ](*t*) using the backward differentiation formula (BDF) along with the time-periodic drivings specified in Eq. 5.

## RESULTS

### Phase diagram, entrainment, and quasi-periodic orbits

Our calculation of the 2D diagram for the PFK model under external periodic driving is summarized in Figs. 2 and 3. Fig. 2A depicts the Lyapunov exponent evaluated for dynamic time trace as a function of the parameters, Ω and *ε* that vary in the ranges of 0 ≤ Ω ≤ 5 and 0≤ *ε* ≤ 1.2*ν*_0_ (see Method for the detail of evaluating Lyapunov exponent). Here, the Ω is the ratio of the system’s natural frequency of oscillation to the frequency of external driving, i.e., Ω = *ω*_*o*_*/ω*_ext_ = *T*_ext_*/T*_*o*_. In Fig. 2B, the period of entrained dynamics *T*_obs_ is calculated on each grid point, if *T*_obs_ is resolved within our simulation time (i.e., *T*_obs_ *< T*_*s*_ = 10^4^ × *T*_ext_); and no specific value of *T*_obs_ is assigned to the grid point if the dynamics is either quasi-periodic or chaotic (see Eq. 25 in Methods). Some of the time traces at their stationary state are plotted in Fig. 2C. The trajectories, marked with star symbols in the diagram of *λ* in Fig. 2A and [*S*](*t*) and [*P* ](*t*) in Fig. 2C, are:

- (red) Frequency-locked orbit entrained to *T*_obs_ ≈ 389 sec generated with *ε* = 0.6 *ν* _0_ and Ω ≈ 1 (*T*_ext_ = *T*_*o*_ ≈ 389 sec).
- (green) Frequency-locked orbit entrained to *T*_obs_ ≈ 389 sec generated with *ε*= 0.4*ν*_0_ and Ω ≈ 1*/*3 (*T*_ext_ ≈ 129 sec, *T*_*o*_ ≈ 389 sec).
- (cyan) *Effectively* quasi-periodic orbit generated with *ε* = 0.3*ν*_0_ and Ω = 2.797.
- (yellow) Chaotic orbit generated with *ε* = 0.92*ν*_0_ and Ω = 2.393.

**FIG. 2.**
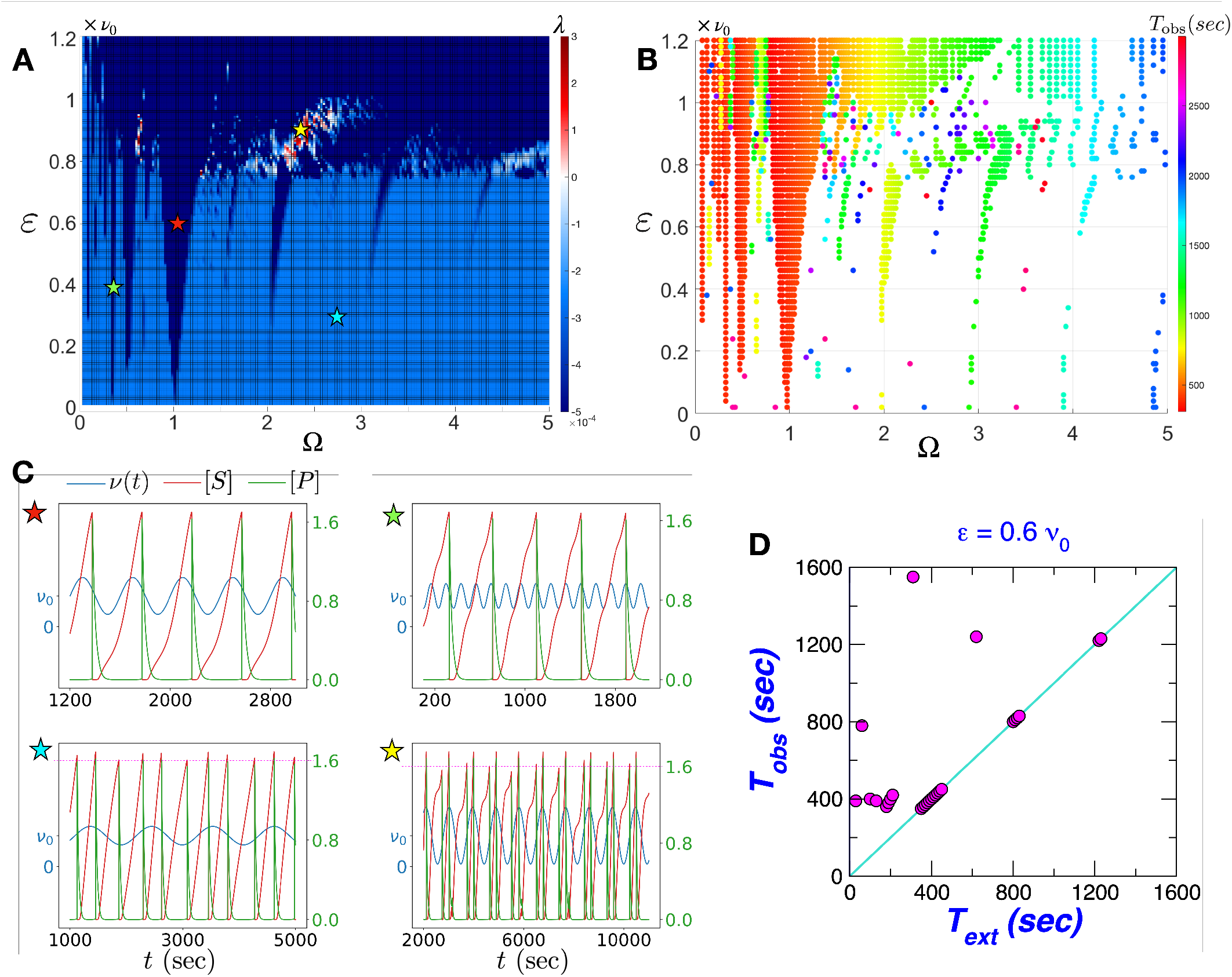
The 2D diagrams of (A) the Lyapunov exponent *λ* and (B) the observed oscillation period *T*_obs_ within the entrainment bands as a function of *ε* and Ω. The oscillation periods less than 3000 sec, which imply that systems are entrained into a frequency-locked state, are depicted with filled circles in rainbow color. (C) The actual time trajectories at parameter values marked with star symbols in different colors. Substrate concentration [*S*](*t*) in red and product concentration [*P* ](*t*) in green in response to periodic driving *ν* (*t*) in blue. The dotted lines in magenta on the panels marked with cyan and yellow stars are drawn to highlight the irregularity of spikes in each interval of the periodic driving (*ν* (*t*)). The trajectories generated at different sets of parameters, *ε* and Ω, marked with star symbols, are: (red) Frequency-locked state entrained to *T*_obs_ *≈* 400 sec (*ε* = 0.6*ν*_0_, Ω ≈ 1); (green) Frequency-locked state entrained to *T*_obs_ ≈ 400 sec (*ε* = 0.4*ν*_0_, Ω ≈ 1*/*3); (cyan) *Effectively* quasi-periodic state (*ε* = 0.3*ν*_0_, Ω = 2.797); (yellow) Chaotic state (*ε* = 0.92*ν*_0_ and Ω = 2.391). (D) The entrainment period (*T*_obs_) as a function of the period of external driving (*T*_ext_) calculated at *ε* = 0.6*ν*_0_. The clusters of points at *T*_obs_ ≈ 400 sec correspond to the periods of entrainment calculated in the Arnold tongues of subharmonic (*T*_ext_ *≈ T*_*o*_*/*2, *T*_*o*_*/*3, *T*_*o*_*/*4) and harmonic (*T*_ext_*≈ T*_*o*_) entrainments. The clusters of points at *T*_obs_ ≈ 800 sec and *T*_obs_ ≈ 1200 sec are due to the Arnold tongues formed under the super-harmonic entrainments. The cyan line corresponds to the condition of *T*_obs_ = *T*_ext_.

**FIG. 3.**
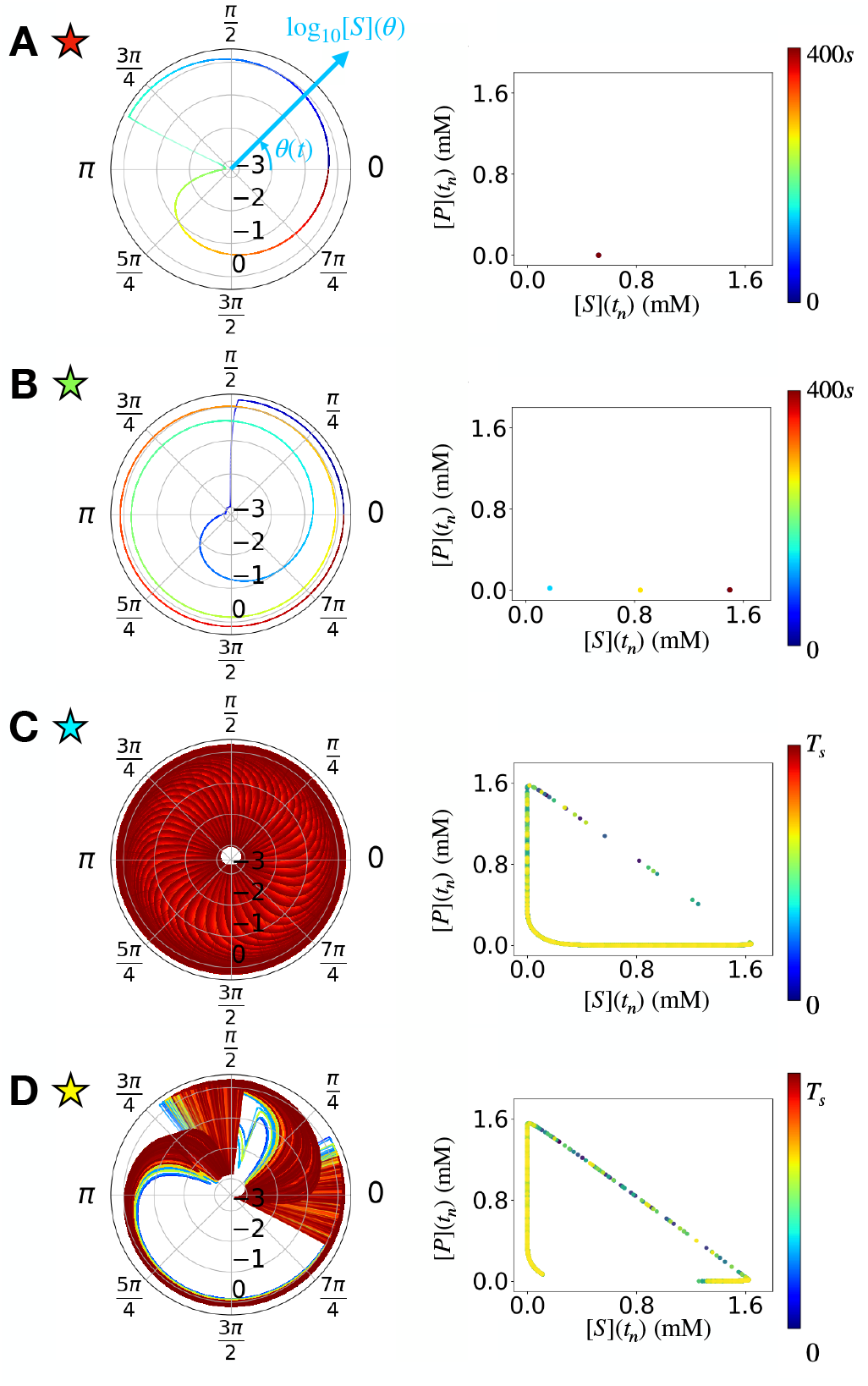
Analysis of the trajectories at different parameter values, marked with star symbols in red (A), green (B), cyan (C), and yellow (D) in Fig. 2A and C. Steady state trajectory of [*S*](*θ*) (radial direction in logarithmic scale) as a function of phase angle *θ* = *ω*_ext_*t* mod 2*π* depicted on the polar coordinate (left), the intersections of trajectory across the Poincaré section ([*S*](*t*_*n*_), [*P* ](*t*_*n*_)) at *t*_*n*_ satisfying *θ*(*t*_*n*_) mod 2*π* = 0 (right). The trajectories and the intersections across the Poincaré section are color-coded in time over 0 *≤ t≤ T*_obs_ for frequency locked states and for simulation time (0*≤ t ≤T*_*s*_) for quasi-periodic and chaotic states.

In the limit of small perturbation (*ε* → 0), if the parameter Ω is rational (*T*_ext_ is commensurate with *T*_*o*_), the system is entrained into a frequency-locked state with a period *T*_obs_, which corresponds to the least common multiple (LCM) of *T*_ext_ and *T*_*o*_, i.e., *T*_obs_ = LCM(*T*_ext_, *T*_*o*_). In contrast, if the ratio of frequencies is irrational (*T*_ext_ is incommensurate with *T*_*o*_), the resulting dynamics is quasi-periodic to display an orbit with an indefinite period (*T*_obs_→ ∞). In practice, if *T*_obs_ = LCM(*T*_ext_, *T*_*o*_) acquired from a rational Ω were much greater such that *T*_obs_ ≫ *T*_ext_, the corresponding frequency-locked state would display an oscillatory orbit with a long period. The corresponding trajectory densely fills the state space and is effectively indistinguishable from the dynamics of a truly quasi-periodic state generated from an irrational Ω. In such a case, we regard the trajectory *effectively quasi-periodic*.

Difference between periodic and (effectively) quasi-periodic trajectories is more clearly grasped by plotting dynamic time traces on a polar coordinate and their piercings on a surface of the Poincaré section (Fig. 3). Dynamic trajectories of frequency-locked states with Ω ≈ 1 (*T*_ext_ ≈ 389 sec) and Ω ≈ 1*/*3 (*T*_ext_ ≈ 129 sec), plotted as a function of phase variable *θ*(*t*) = *ω*_ext_*t* mod 2*π* with *ω*_ext_ = 2*π/T*_ext_ on the polar coordinate, form closed loops with a single (Fig. 3A, left) and three revolutions (Fig. 3B, left), displaying a single (Fig. 3A, right) and three piercings (Fig. 3B, right) on the Poincaré section at *θ*(*t*_*n*_) = 0, respectively. In comparison, the trajectory of an effectively quasi-periodic state with Ω = 2.797 (*T*_ext_ = 1085 sec, the trajectory marked with cyan star in Figs. 2 and 3), which results in an orbit with *T*_ext_ ≈ 420980 sec, almost fully covers the phase space on the polar coordinate as well as gives rise to the continuous piercings on the corresponding Poincaré section.

A finite amplitude (*ε* ≠ 0) coupled to the oscillatory mode of the system expands the parameter range of Ω for the frequency-locked state, and thus widens the entrainment bands, forming the Arnold tongues [33–36] (Fig. 2B). Consistent with the experiment by Boiteux *et al*. [4], the subharmonic entrainments to *T*_obs_ = *nT*_ext_ ≈ *T*_*o*_ ≈ 389 s for Ω(= *T*_ext_*/T*_*o*_) ∼1*/n* with *n* = 2, 3, 4, · · · are confirmed for Ω ≲ 1 (Fig. 2B). For Ω *>* 1, we observe super-harmonic entrainments to *T*_obs_ ≈ 1200 s at Ω ≈ 3*/*2, *T*_obs_ ≈ 800 s at Ω ≈ 2, *T*_obs_ ≈ 1200 s at Ω ≈ 3, and *T*_obs_ ≈ 1600 s at Ω≈ 4 (Fig. 2B). The width of the Arnold tongue is the greatest for the harmonic entrainment, i.e., Ω ∼ 1.

In principle, unless *λ >* 0, all the time traces generated at each grid point of rational Ω (Fig. 2) are still characterized with a finite periodicity *T*_obs_ *<* ∞ . However, outside the entrainment bands, the 2D phase diagram is in fact densely populated by quasi-periodic states since the density of irrational Ω is significantly higher than that of rational Ω. The Lebesgue measure of irrational Ω, giving rise to quasi-periodic orbits, is 1 for *ε*→ 0 [17, 18]. Meanwhile, an increasing value of *ε* leads to expansions of the entrainment bands, which in turn reduces the region associated with the quasi-periodicity. Fig. 2D, which visualizes the periods of entrainment (*T*_obs_) for a given period of external driving (*T*_ext_) at *ε* = 0.6*ν*_0_, hints at the winding number [34, 35] of the map associated with our model.

### Chaotic state

The conditions that give rise to a chaotic orbit are of great interest. The trajectories of [*S*](*t*) and [*P* ](*t*) at a chaotic state (*ε* = 0.92*ν*_0_, Ω = 2.391) which display a train of spikes with irregular height (Fig. 2C, yellow star) are similar to those with quasi-periodicity (Fig. 2C, cyan star), and the patterns of piercings on the Poincaré section are also qualitatively similar (Figs. 3C and D). However, together with the sign of Lyapunov exponent, chaotic orbit is differentiated from quasi-periodic orbit when the trajectory is drawn on the polar coordinate of phase angle. The phase space represented in the polar coordinate (Fig. 3) is only partly filled with the chaotic orbit, whereas it is fully filled with the quasi-periodic orbit.

The chaotic states with positive Lyapunov exponents (*λ >* 0) are sporadically identified in the phase diagram. In Fig. 2A, they are localized in the vicinity of Ω ≈ 0.5 and Ω≈ (2 −2.5) for *ε* ≈ (0.8 −1)*ν*_0_. Fig. 4A extends the phase diagram displayed in Fig. 2A to a wider range of *ε*. It is clear from Fig. 4A that while chaotic states are scarce under strong driving (*ε* ≳ *ν*_0_), they are still found at *ε* ≈ 4*ν*_0_ and 7*ν*_0_ in the range of 0 *<* Ω ≲ 0.3. The inset of Fig. 4A magnifies the region of phase diagram where chaotic states are frequently observed, and the chaotic maps at fixed Ω = 0.257 (Fig. 4B) visualize qualitative changes of the periodic orbits, [*S*](*t*) and [*P* ](*t*) with increasing *ε*. The periodic orbits generated at four specific values of *ε* marked with the arrows in different colors in the inset of Fig. 4A and Fig. 4B are shown in Fig. 5. Straightforwardly gleaned from the number of piercings in the Poincaré section (the panels in the rightmost column of Fig. 5), the system is in the frequency-locked states of period-3 (*ε* = 5*ν*_0_), period-5 (*ε* = 6*ν*_0_), and period-2 cycles (*ε* = 8*ν*_0_); however, it displays a chaotic behavior at *ε* = 7*ν*_0_. The irregularities in the size and the number of spikes of the chaotic orbit (*ε* = 0.7*ν*_0_) are seen in the leftmost panel in Fig. 5, and the other remaining panels of Fig. 5 capture the characteristic features of chaotic states.

**FIG. 4.**
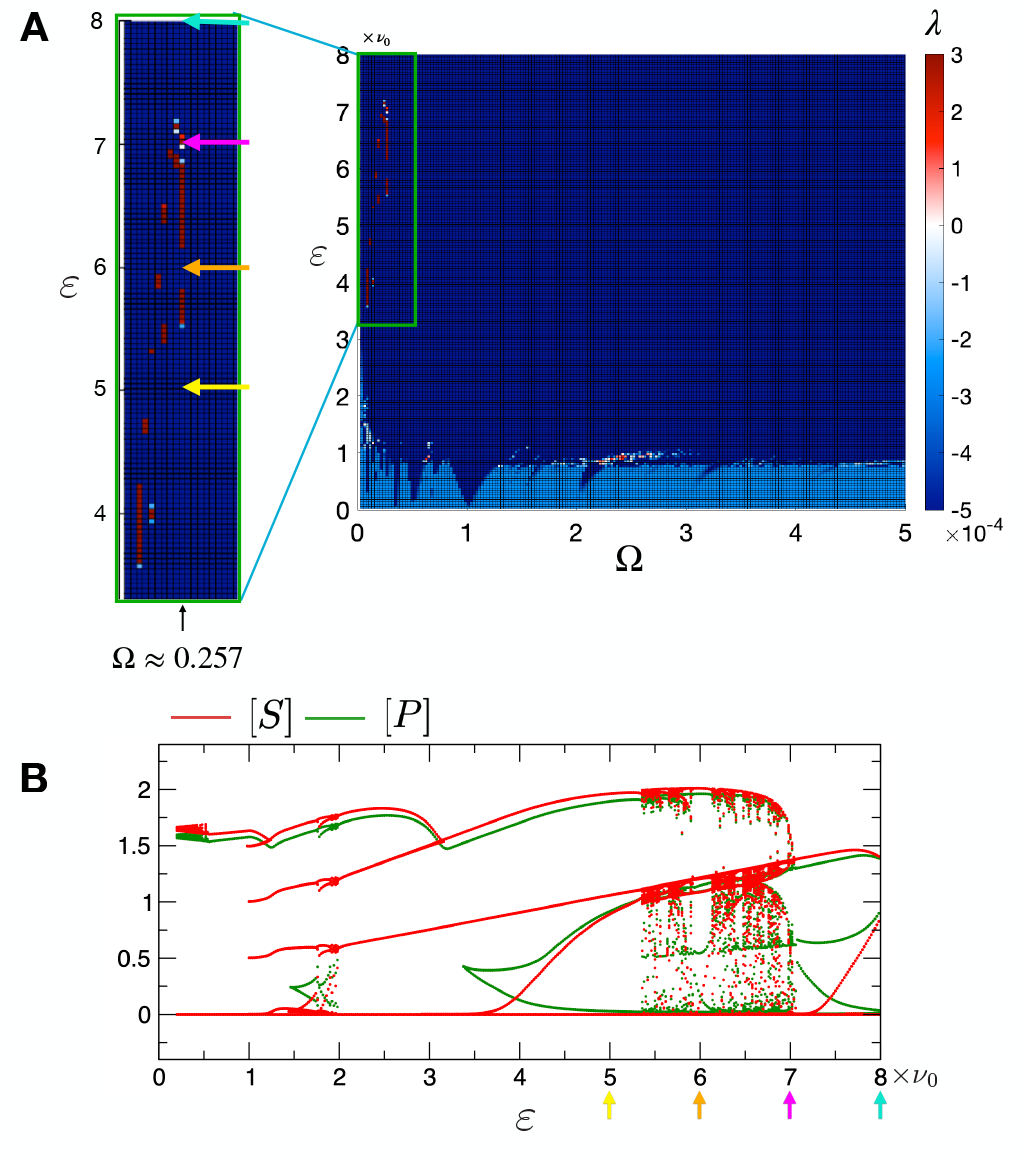
Transition to chaos demonstrated for *ε* extended to large values at fixed Ω(= *T*_ext_*/T*_*o*_ = 100*/*389) ≈ 0.257. (A) Phase diagram of the Lyapunov exponent *λ* extended to large *ε*. The inset on the left enlarges the region of phase diagram with chaotic states. The trajectories of [*S*](*t*) and [*P* ](*t*) at four distinct sets of parameter values, indicated by the arrows, are demonstrated in Fig. 5. (B) Chaotic maps for [*S*](*t*) and [*P* ](*t*) as a function of *ε* calculated at fixed Ω ≈ 0.257.

**FIG. 5.**
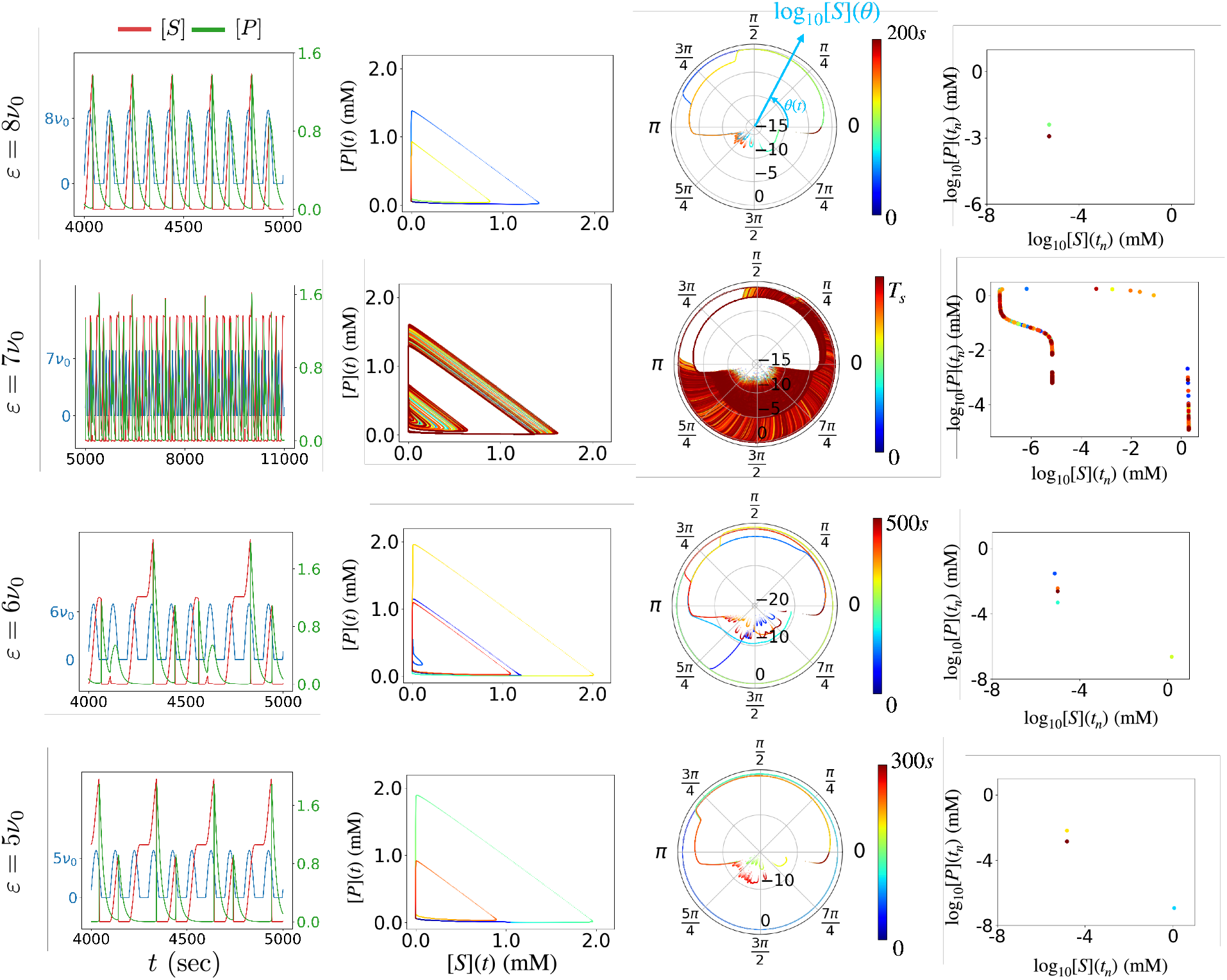
Trajectories generated under the conditions marked with four arrows in different colors in Fig. 4. The dynamical state that exhibits a period-3 cycle at *ε* = 5*ν*_0_ changes to a chaotic state at *ε* = 7*ν*_0_, and to a state with period-2 cycle at *ε* = 8*ν*_0_. The time-evolutions of [*S*](*t*) (red) and [*P* ](*t*) (green) generated in response to *ν* (*t*) (blue) on the first column, and the plots of [*S*](*t*) versus [*P* ](*t*) are shown on the second column. On the third column, the steady state trajectory of substrate concentration [*S*](*θ*) is depicted as a function of phase angle *θ* = *ω*_ext_*t* mod 2*π* on the polar coordinate. The rightmost panels show the intersections of dynamic time trace across the Poincaré section.

### Mapping onto the circle map

To analyze the model of glycolytic oscillation under the periodic driving more systematically, we recast the original set of ODEs in the following form:

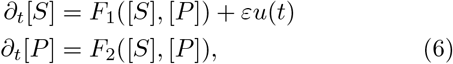

where *u*(*t*) = sin (*ω*_ext_*t*) = *u*(*t* + *T*_ext_) is a *T*_ext_-periodic perturbation term which can be defined consistently with Eq.5. For *ε* = 0, the system is left unperturbed, exhibiting a stable limit cycle dynamics on **x**(*t*) = ([*S*], [*P* ]) with the oscillation frequency *ω*_*o*_(= 2*π/T*_*o*_) (Fig. 1C).

For *ν*_0_ = 0.005 mM·*s*^−1^, *k*_*d*_ = 0.05 *s*^−1^, and *ε* = 0 (the yellow star in the phase diagram depicted in Fig. 1B), the substrate and product concentrations ([*S*](*t*), [*P* ](*t*)) undergo oscillation around the (unstable) fixed point ([*S*^*^], [*P* ^*^]) = (0.22mM, 0.10mM) (see the phase portrait in Fig. 1C), and this dynamics can be described using time-dependent phase angle and amplitude defined as *θ*(*t*) = tan^−1^(*δ*[*P* ](*t*)*/δ*[*S*](*t*)) ∈ [0, 2*π*) and 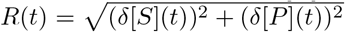, where *δ*[*S*] = [*S*] − [*S*^*^] and *δ*[*P* ] = [*P* ] − [*P* ^*^]. While the phase angle *θ*(*t*) is 2*π*-periodic, its variation is still non-uniform in time, i.e., *dθ*(*t*)*/dt* ≠ *constant*, as suggested by the varying size of velocity field in Fig. 1C. Thus, one can consider uniformizing the phase angle by defining a new phase angle *ϕ* (*θ*),

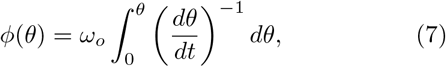

which changes linearly in time for a limit cycle dynamics and satisfies *ϕ* (*θ* + 2*π*) = *ϕ*(*θ*) + 2*π*.

Under a *T*_ext_-periodic driving, *u*(*t* + *T*_ext_) = *u*(*t*) with small *ε*, one can formally recast Eq. 6 to an equation of motion in continuous time for the phase angle *ϕ*.

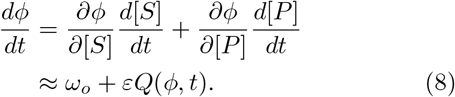

Where 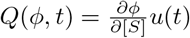. Integrated over a single period of driving *T*_ext_, i.e., from *t*_*n*_ to *t*_*n*+1_, where *t*_*n*_ = 2*πn/ω*_ext_, Eq.8 can be written as a one-dimensional mapping from *ϕ*_*n*_ to *ϕ*_*n*+1_, which corresponds to the *circle map* with an additional perturbation,

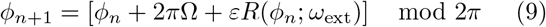

where *ϕ*_*n*_ = *ϕ* (*t*_*n*_), Ω = *ω*_*o*_*/ω*_ext_ = *T*_ext_*/T*_*o*_, and 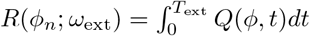.

As *Q*(*ϕ, t*) is a 2*π*-periodic function of *ϕ* and a *T*_ext_-periodic function of time *t*, the phase space of Eq. 8 is defined on a 2D torus of 0 ≤ *ϕ <* 2*π* and 0 ≤ *t < T*_ext_. For *ε* = 0, the dynamics of the uniformized phase angle forms a limit cycle with frequency *ω*_*o*_, satisfying *ϕ* (*t*) = *ω*_*o*_*t* + *ϕ*_0_. Importantly, there exists a one-to-one correspondence between the points on the unperturbed trajectory **x**_*o*_(*t*) = ([*S*](*t*), [*P* ](*t*)) and the phase angle *ϕ* (*t*), such that **x**_*o*_ = **x**_*o*_(*ϕ*) is an invertible function of *ϕ*. A forced dynamical system with *ε* /= 0 still sustains its oscillatory trajectory **x**_*ε*_(*ϕ*) around the stable limit cycle **x**_*o*_(*ϕ*) as long as *ε* remains small. However, when *ε* is too large, **x**_*ε*_ = **x**_*ε*_(*ϕ*) becomes non-invertible, so that the one-to-one mapping between **x**_*ε*_ and *ϕ* no longer holds, which leads to disruption of the torus [37]. In the case of the *sine circle map, R*(*ϕ*_*n*_) in Eq. 9 is given as *R*(*ϕ*_*n*_) = sin *ϕ*_*n*_, and hence the map becomes non-invertible if 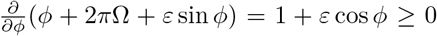 for some *ϕ*, which is realized when *ε* ≥ 1. In our PFK model of glycolytic oscillation, it is found that the transition to non-invertibility, which permits the formation of chaotic orbits, is made for *ε* ≳0.8*ν*_0_ (Fig. 2A).

## DISCUSSIONS

The actual glycolytic oscillation is far more complicated than the simplified model explored here [8, 11, 12, 38–40]. However, the overall dynamical responses of the system to the external periodic driving, are recapitulated as (i) the harmonic and sub/super-harmonic entrainments inside the Arnold tongues at weak-to-moderate driving, (ii) quasi-periodic states outside the Arnold tongues, and (iii) emergence of chaotic behaviors at strong driving. These are generic features expected for a dynamical system demonstrating a self-sustained oscillation [36] and should neither be limited to the model of glycolytic oscillation nor depend on its microscopic details. Despite the diversity of network designs for biochemical oscillations, one can still utilize the general notions of dynamical system to gain reasonable understanding of their responses to periodic drivings.

Whereas this study discusses the emergence of chaotic behaviors from the conventional PFK model of glycolysis under *non-autonomous external driving, chaotic dynamics can also arise autonomously for a more complex version of glycolysis. Goldbeter and coworkers [54] have studied another PFK model of glycolysis with three variables constituting two positive feedback loops, showing that the model, albeit autonomous, exhibits chaotic dynamics as well as both simple and birhythmic oscillations*.

In order for a yeast cell to display glycolytic oscillations, the glucose injection rate along with the product degradation rate must be in a proper range. The phase diagram of PFK model built under the condition of constant glucose injection rate (Fig. 1B) suggests that an increased glucose injection rate surpassing *ν*_0_ ≈ 0.01 mM/s (*k* = 0.05 *s*^−1^) results in the cessation of glycolytic oscillation, which could imply a physiological state potentially associated with the hyperglycemia. This study, however, suggests that the dynamical response of the system exhibiting a limit cycle to the external forcing is extremely rich. As long as the glucose is injected in a time-varying (periodic) manner over a proper range of frequency and amplitude, the system can be entrained into the Arnold tongues, still sustaining its oscillatory dynamics around its natural frequency. Although the role of glycolytic oscillations in physiology is not fully elucidated, glycolytic oscillations have been suggested as a pacemaker of other physiological oscillations [41–43]. Specifically, an oscillation of NADH, a product of glycolysis and the precursor of Kreb/electron transfer cycle, drives the oscillation of mitochondrial membrane potential [44]. Sustained oscillations in lactate released from islets of Langerhans provide a mechanism for pulsatile insulin secretion from *β*-cells [5]. Furthermore, acetaldehyde, a membrane-permeating metabolite from glycolysis, mediates cell-to-cell communication among a yeast cell population leading to in-phase oscillations [45–49].

The enhancement of the harmonic and sub/super-harmonic entrainments of glycolytic oscillation under the weak-to-medium periodic drivings (*broadening of the entrainment bands*) could be discussed from a more general perspective considering a phase angle dynamics under non-autonomous time-varying perturbation expressed in the form of Eq.8. Specifically, recent studies on the synchronization among phase oscillators coupled with a coupling strength *γ* that can be mapped to *ε* in the present study, have also shown enlarged Arnold tongues with increasing *γ* [50], indicating the enhanced stability of oscillatory dynamics or *chronotaxicity*, in other words the ability of a self-sustained oscillator to resist non-autonomous perturbations. Along with these studies [50, 51], our study substantiates that the glycolytic oscillation of a single cell can indeed be influenced by the oscillatory metabolic signals from the surrounding cells, and explains the ease of inter-cellular synchronization and enhanced stability of oscillations in cell populations [49, 52, 53]. However, an excessive amount of glucose injection can bring the system into disarray, rendering the dynamical state of the system chaotic, in other words, hyper-sensitive to initial conditions and unpredictable.

## METHODS

### Evaluation of Lyapunov exponents

For general *d*-dimensional continuous-time dynamical equation satisfying a set of ODEs with the vector function **F**[**x**(*t*)] over the *d*-dimensional vector **x**(*t*),

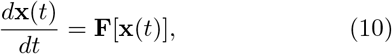

its variation of variables, **x** → **x** + *δ***x**, yields

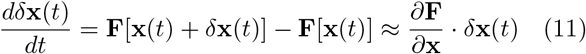

for *δ***x**(*t*) → 0, and it formally satisfies

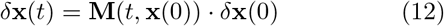

where 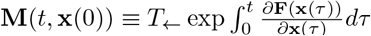 denotes an *d*×*d* time-ordered exponential operator. The Lyapunov exponent of a dynamical system dictates the growth rate of the size of the variation. Thus, one can consider

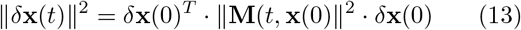

with

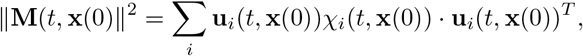

where **u**_*i*_(*t*, **x**(0)) and *χ*_*i*_(*t*, **x**(0)) are the *i*-th eigenvector and eigenvalue of ||**M**(*t*, **x**(0)) || ^2^ matrix, respectively. The Lyapunov exponent associated with the change in the vector size ||*δ***x**(*t*) ||∼ *e*^*λt*^ ||*δ***x**(0) || is calculated as follows [55].

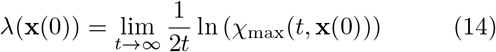

where *χ*_max_ is the largest eigenvalue among the set of eigenvalues {*χ*_*i*_} of ||**M**(*t*, **x**(0)) || ^2^ matrix.

For a practical numerical procedure to evaluate the Lyapunov exponent, we used the one proposed by Benettin *et al*. [56]. We consider *d* possible variations from *d*-dimensional vector **x**(0), say, *δ***x**_1_(0), *δ***x**_2_(0) …, *δ***x**_*d*_(0). Since these variations are not generally orthonormal to each other, the Gram-Schmidt procedure is used to obtain a corresponding set of orthonormal vectors 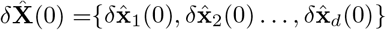, which span the same sub-space by *δ***X**(0) = {*δ***x**_1_(0), *δ***x**_2_(0) …, *δ***x**_*d*_(0) } . Specifically, we build mutually orthogonal vectors

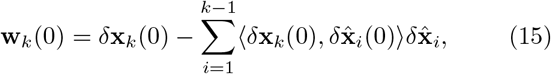

and normalize them using

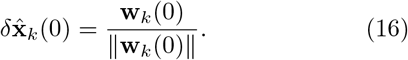

We integrate Eq. 10 to calculate the next position **x**(*τ* ) and employ Eq. 12 to calculate the next variations *δ***X**(*τ* ) ≡ [*δ***x**_1_(*τ* ), *δ***x**_2_(*τ* ) …, *δ***x**_*d*_(*τ* )]. We then again apply the Gram-Schmidt procedure to obtain the corresponding set of orthonormal vectors 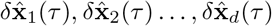. We repeat this procedure of integration and orthonormalization over the time duration of *Nτ* for the full time trace.

At time *τ* ≫ 1, we expect

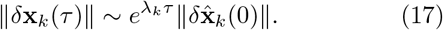

Hence, the Lyapunov exponent corresponding to the growth rate of the variation in the *k*-th direction is given by

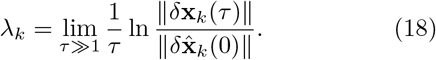

This can be cast into a matrix form

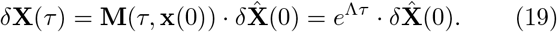

where **M**(*τ*, **x**(0)), *δ***X**(0), and Λ = (*λ*_1_, …, *λ*_*d*_) · 𝕀 are *d* × *d* matrices, and it yields

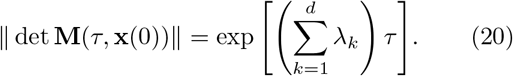

Therefore,

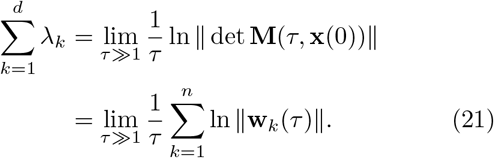

Note that the transformation 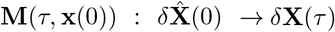 maps the *d*-dimensional *hypercube* of volume 1 formed by the *d* orthonormal vectors of variation 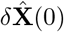 into an *d*-dimensional *parallelopiped* made of *δ***X**(*τ* ). Thus, (i) 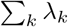 measures the growth rate of the *d*-dimensional volume from the hypercube to the parallelopiped. (ii) The second line of the Eq.21 follows from the fact that the determinant of the operator **M**(*τ*, **x**(0)) is identical to the volume of the *d*-dimensional parallelopiped, Vol(*P* ), and this volume can be calculated using the product of mutually orthogonalized vectors of *δ***x**_*k*_(*τ* ), i.e., **w**_*k*_(*τ* ) (Eq.15). Taken together,

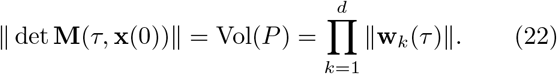

In our study, we aim to decide the growth (or convergence) rate of the trajectory averaged over the period of the external driving along the *k*-th variation. Provided that the time traces are generated over *N* iterations of the period at steady state, we set *τ* = *T*_ext_ and consider calculating the Lyapunov exponent averaged over the *N* times of driving period.

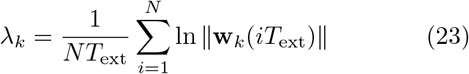

Finally, the Lyapunov exponent characterizing the dynamical behavior of the system is obtained from the set of {*λ*_1_, *λ*_2_ …, *λ*_*d*_} with *λ*_1_ *> λ*_2_ *>* … *> λ*_*d*_, and the largest Lyapunov exponent *λ*_1_ decides the dynamical behavior of the system at a long time limit.

Our system subject to an external periodic driving (Eq. 6) can be recast in the form of a set of ODEs, 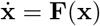, with three variables (*d* = 3), **x** = ([*S*], [*P* ], *ψ*),

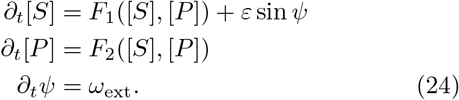

Since *ψ* is *T*_ext_-periodic, the corresponding Lyapunov exponent is always 0. Hence, *λ*_1_ = 0 and *λ*_2_, *λ*_3_ *<* 0 for the case of periodic or quasi-periodic orbits, whereas *λ*_1_ *>* 0 for chaotic orbits. In the phase diagrams of Lyapunov exponents in Figs. 2A and 4A, we report the value of exponent *λ*, such that we use the positive largest Lyapunov exponent (*λ* = *λ*_1_ *>* 0) to quantify the growth rate of variation for chaotic states and the negative largest Lyapunov exponent (*λ* = *λ*_2_ *<* 0) to quantify the convergence rate for periodic or quasi-periodic states:

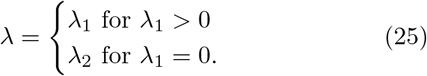

## ACKNOWLEDGEMENTS

We thank Dr. Won Kyu Kim and Dr. Andrus Giraldo for careful reading of the manuscript and helpful comments. This work was supported by KIAS Individual Grants, CG076501 (P.K.) and CG035003 (C.H.) at Korea Institute for Advanced Study. We thank the Center for Advanced Computation in KIAS for providing computing resources. The computer codes for the numerical integration of ODEs and the evaluation of Lyapunov exponents can be accessed at the Github repository (https://github.com/purn25/Glycolytic_oscillations).

## APPENDIX

### Effect of stochastic forcing on self-sustained oscillations

In comparison with the periodic forcing (Eq. 5), the stochastic forcing can be written as *ν* (*t*) = *ν* _0_ + *ξ* (*t*) with ⟨ *ξ* (*t*) ⟩ = 0 and ⟨ *ξ* (*t*) *ξ* (*τ* ) ⟩ = *εδ*(*t*−*τ* ). Boiteux *et al*. varied the glucose injection rate (30 – 180 mM/hr ≃ 8.33 × 10^−3^ − 0.05 mM/sec) “stochastically” around its average value *ν*_0_(= 0.005 mM/sec), finding that the system retains its autonomous oscillation around its natural frequency [4].

In the light of Eq. 8, their finding can be formulated into the following dynamics of a limit cycle in a phase variable *ϕ*: *d ϕ /dt* = *ω*_0_ + *ξ*(*t*). Thus, it follows that *µ*_*ϕ*_ ≡ ⟨ *δϕ*(*t*) ⟩ = *ω*_0_*t* and 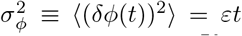, where ⟨… ⟩ denotes the average over trajectories. If one were to have a relative error (noise-to-signal ratio) less than 1, i.e., 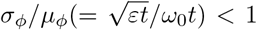, one obtains a condition Of 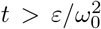. This means that even when subject to a stochastic forcing with a strength *ε*, the underlying phase variable dynamics of a self-sustained oscillation with the frequency *ω*_0_ can still be read out as long as the observation time *t* is greater than 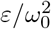.

